# Dynamics of lipids in the yeast lipid droplets

**DOI:** 10.1101/2021.10.22.465485

**Authors:** Margarita Fomina, Eugene Mamontov, Hugh O’Neill, Dan Close, Jose M. Borreguero, Jennifer Morrell-Falvey

## Abstract

The physical properties and chemical composition of lipid droplets inside cells of the yeast *Cryptococcus curvatus* were investigated using quasi-elastic neutron scattering and mass spectrometry with complementary surface modeling using 3D microscopy. With temperature decrease from 310 to 280 K, their phase remained fluid, i.e., the droplets remained in the physiological state, unlike synthetic lipid membranes that transition to a gel phase. The lipid dynamics in the droplet was described by a model implying diffusion of the lipid and its hydrocarbon chains. The diffusion coefficient of the lipid chains (274×10^−3^ Å^2^/ps at 310 K) was much higher than that observed in a synthetic lipid membrane because of the larger volume (up to 12 Å) for the local dynamics in the droplet. These physical properties were correlated with the types of lipids composing the droplet. Based on that, the lipid packing and resulting energetic value of the yeast droplets are discussed in relation to their usefulness as biofuels.

---

Every organism stores lipids as an energy resource for metabolic processes. Some unicellular eukaryotes, such as oleaginous yeasts, synthesize lipid reserves *de novo* when cultivated in a growth medium containing an excess of carbon over nitrogen [1,2]. This lipid production occurs especially during the late-growth stage of the cells when nutrients essential for proliferation—nitrogen, sulfate, and phosphorus—become scarce and excessive carbon is redirected toward lipogenesis [1,3–5], producing primarily triacylglycerol (TAG) and steryl ester [6,7]. A TAG is formed when three fatty acids (a hydrocarbon chain with carboxyl group) are joined together by a glycerol molecule, and a steryl ester is produced when a fatty acid attaches to a sterol. It is well accepted that these lipids are collected between the membrane bilayers of the endoplasmic reticulum (ER) having the enzymes for lipid synthesis [8]. The synthesized lipids are nonpolar molecules, forming a hydrophobic core, which tears off the ER, taking a phospholipid monolayer of the membrane with the associated proteins as a shell for its cytoplasmic environment. A lipid droplet (LD) thus formed becomes an individual cellular organelle (Fig. 1).

**FIG. 1.**
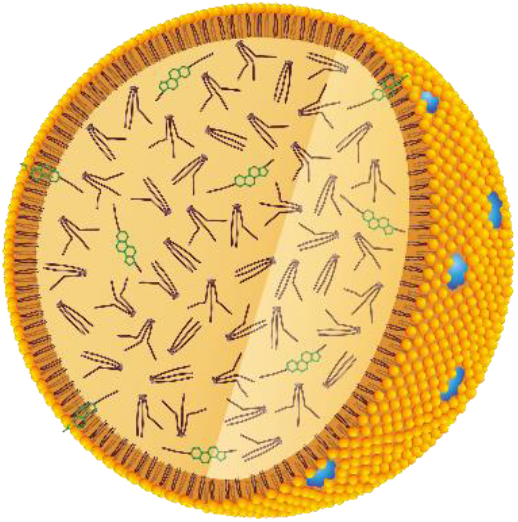
A sketch of the yeast LD in cross-section. The TAGs (violet) and steryl esters (green/violet) are in the core, surrounded by a monolayer of phospholipids (orange/violet) with embedded proteins (blue).

From the applied side, the LDs of microorganisms are a source of hydrocarbons for the use in food, animal feed, precursors for chemical synthesis, and biofuels [9–11]. Considering the diminishing supplies of nonrenewable energy sources, particularly oleaginous yeasts represent a source of renewable and environmentally-safe hydrocarbon fuel owing to their higher genetic tractability relative to algal counterparts, faster growth rate relative to bacterial analogs [12], and ability to grow on low-cost substrates [13–19].

Revealing the properties of LDs will allow to optimize them as a source of biofuels. We explored intrinsic physical properties of lipids in the droplets, probing them in intact yeast cells. The lipid dynamics at different temperatures was revealed by quasi-elastic neutron scattering (QENS) at a high-resolution near-backscattering spectrometer [20]. QENS resolves molecular motions on a time scale from a few to hundreds of picoseconds and a length scale from a few to tens of Ångstroms. Neutrons represent a suitable probe for the dynamics of hydrogen-rich molecules such as lipids due to the large incoherent scattering cross-section of hydrogen [21]. Note that dynamics of in-droplet lipids was resolved; due to the LD’s large size, its center-of-mass diffusion is too slow to be resolved by QENS.

A reference for our investigation was a previous QENS study of vesicles made of a unilamellar bilayer of DMPCs (1,2-dimyristoyl-sn-glycero-3-phosphocholines) [22,23]. The amphiphilic and saturated DMPCs pack uniformly in a bilayer. The yeast droplet core contains hydrophobic lipids, which might pack randomly, and that would be reflected in their dynamics.

The present work advances lipid research by studying properties of lipids in their native environment. Here, the LD host was the oleaginous yeast *Cryptococcus curvatus* ATCC20509, accumulating LDs up to 60% of the cell volume [24]. The sample preparation is reported in the Supplemental Material (SM) [25].

The temperature dependence of the elastic incoherent neutron scattering (EINS) from the sample represents a response of the overall dynamical behavior of the sample to a temperature change. EINS continuously decays with temperature increase if there is a diffusion process in the sample. A step-like decrease of the elastic intensity upon heating (or an increase in cooling) is typically associated with a phase transition of the system.

EINS from the yeast LDs (Fig. 2(a)) decreases over the measured temperature range due to a diffusive process revealed by QENS signal hereafter. The absence of a step-like change in the elastic intensity points to one phase of the yeast LDs from 280 to 310 K, in contrast to the DMPC bilayers undergoing a gel-to-fluid transition at 296 K [22]. Thus, the LDs stay either in the gel or fluid phase within this temperature range. The analysis of QENS signal provided below indicated rather a fluid phase of the LDs. The presence of unsaturated lipids and steryl esters as well as the length of lipid chains affects the transition temperature of the lipid system [26]. Using gas chromatography/mass spectrometry (GC/MS) we determined the lipid composition of the yeast droplets. We found an almost equal proportions of unsaturated and saturated lipids [25]. Comparing with the synthetic membrane made of POPC (1-palmitoyl-2-oleoyl-sn-glycero-3-phosphocholine), a phospholipid with saturated and unsaturated hydrocarbon chains, undergoing the phase transition at 271 K [27], we can speculate that a gel-to-fluid transition of the LDs occurs below 280 K.

**FIG. 2.**
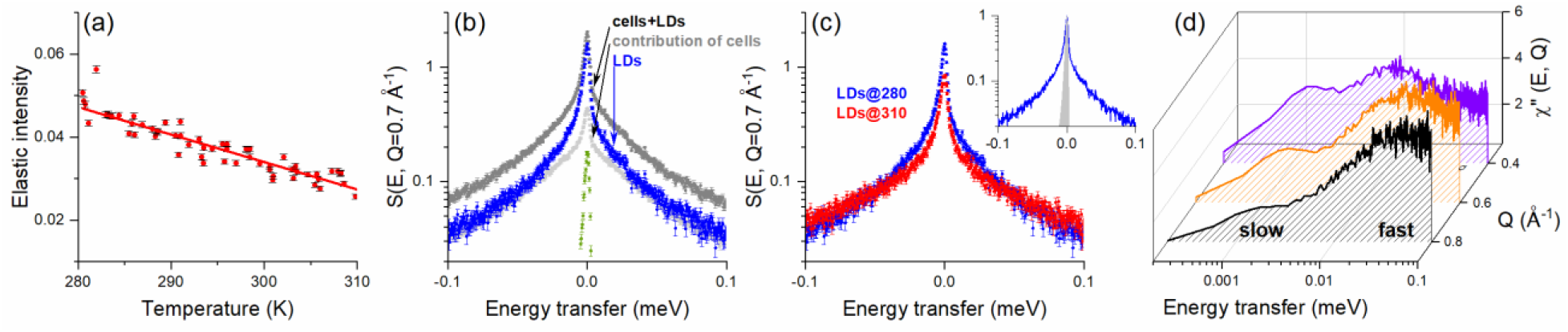
(a) The Q-summed elastic incoherent intensity of the yeast LDs on heating. The errors are standard deviations. Solid line is drawn as a guide to the eye. (b) QENS spectra at 280 K of the cells with LDs (dark gray), contribution of the cells (light gray), empty sample holder (green), and derived LDs signal (blue). (c) QENS signal of LDs at 280 and 310 K; inset: a comparison of peak-normalized LDs signal at 280 K (blue) with the instrument resolution (gray shaded) highlights the accessible energy (time) scale of the lipid dynamics. (d) Dynamic susceptibility of the LDs at 280 K for Q=0.4 Å^-1^ (violet), 0.6 Å^-1^ (orange), 0.8 Å^-1^ (black). The relaxations at longer and shorter times are indicated with the data at 0.8 Å^-1^.

The microscopic dynamics of the LDs was investigated in detail via the QENS signal. Because of the strong incoherent neutron scattering from the protonated lipids, the intensities of the cells having LDs were well above the intensities of the cells (Fig. 2(b)). Executing a 3D surface modeling on the cells with LDs in the IMARIS software, we obtained the volume fractions of 43% for the LD and 57% for its cell [25]. The QENS signal of the LDs was derived following the subtraction of the cell contribution, as explained in the SM [25]. At 310 K the lipids move faster as evidenced by decreased elastic and increased width of QENS contributions in comparison to the spectrum at 280 K (Fig. 2(c)).

The dynamics of lipids can be complex due to different degrees of freedom depending on the lipid organisation. For example, the dynamics of lipids in a bilayer have been modelled by six time-dependent processes [28]. In a QENS experiment distinct lipid motions can be revealed through the relaxation times of its molecules. A spectrum of the dynamic susceptibility *χ*″(*E, Q*) [29] reveals the molecular relaxation via a peak with a maximum corresponding to a characteristic relaxation time and can be calculated from the dynamic structure factor corrected for the Bose occupation number as

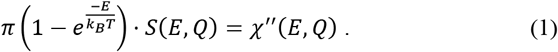

The dynamic susceptibility of the LDs revealed a distinct peak at the energy of ∼10^−3^ meV, followed by a weaker peak at the higher energy of ∼10^−2^ meV (Fig. 2(d)). In terms of a time scale, these relaxations occur at longer and shorter times, respectively. Towards higher Q, i.e. at a smaller length scale, the relaxations occurred faster, which is evidenced by the shift of peaks to higher energies. At higher Q, the contribution of the long-time relaxation decreased resembling a slow long-range translational molecular motion barely detectable on the spatial scale; whereas, the short-time relaxation became dominant in the total dynamics reminiscent of a fast localized molecular motion well detectable on that scale. Similar to the assumption made for the QENS data of the DMPC bilayers [22], we associated the slow motion with the diffusion of whole lipid across the LD core (global diffusion) and the fast motion with the diffusion of bound lipid chains (local diffusion).

For the quantitative analysis of the LDs spectra, we applied the mathematical function used by Sharma et al [22]. The total dynamic structure factor was an analytical convolution of the two structure factors, *S*_*global*_(*E, Q*) and *S*_*local*_(*E, Q*), multiplied by the Debye-Waller factor describing the effect of temperature-dependent atomic displacements on the signal intensity:

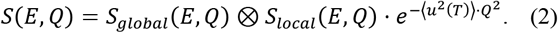

The first structure factor describes the diffusion of whole lipid by the Lorentzian function:

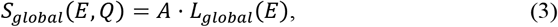

where *A* takes account of the total of H-atoms moving slowly.

The second structure factor describes the diffusion of bound lipid chains. The diffusion of the H-atoms is restricted by the bonds to the carbon chains, and to describe it we applied the neutron incoherent scattering law for diffusion in a potential of spherical symmetry [30]. It is the sum of an elastic component, referred to the elastic incoherent structure factor (EISF) describing the geometry of the limited space where the diffusion occurs, and a quasi-elastic component of the actual motion. Mathematically, they are expressed by a Dirac delta and a normalized Lorentzian function, respectively:

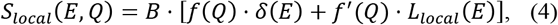

where *f*′(*Q*) *=* 1 − *f*(*Q*) and *B* takes account of the total of H-atoms moving fast.

The normalization of *S*(*E, Q*) to its integral over the energy range suppressed the intensity coefficients *A* and *B*, and the Debye-Waller factor, resulting in

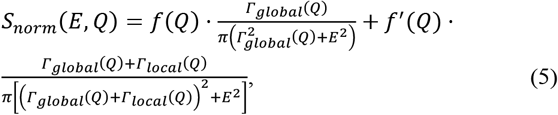

where *Γ*(*Q*) is the half width at half maximum (HWHM) of the Lorentzian function and the normalized *f*(*Q*) is the EISF. The details on normalization of *S*(*E, Q*) are reported in the SM [25].

The function for Q-by-Q fitting of the experimental spectra is a sum of the normalized composite structure factor numerically convoluted with the instrument resolution, and a background constant:

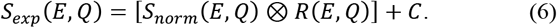

Fig. 3(a) shows the normalized spectra of the yeast LDs fitted by the function *S*_*exp*_(*E, Q*). The constant *C* was determined at Q=0.3 Å^-1^ where the background can be discriminated from the very narrow QENS contribution. In the energy range of ±0.1 meV the background partially accounts for a small contribution of faster processes (vibrations and relaxations of lipid chains).

**FIG. 3.**
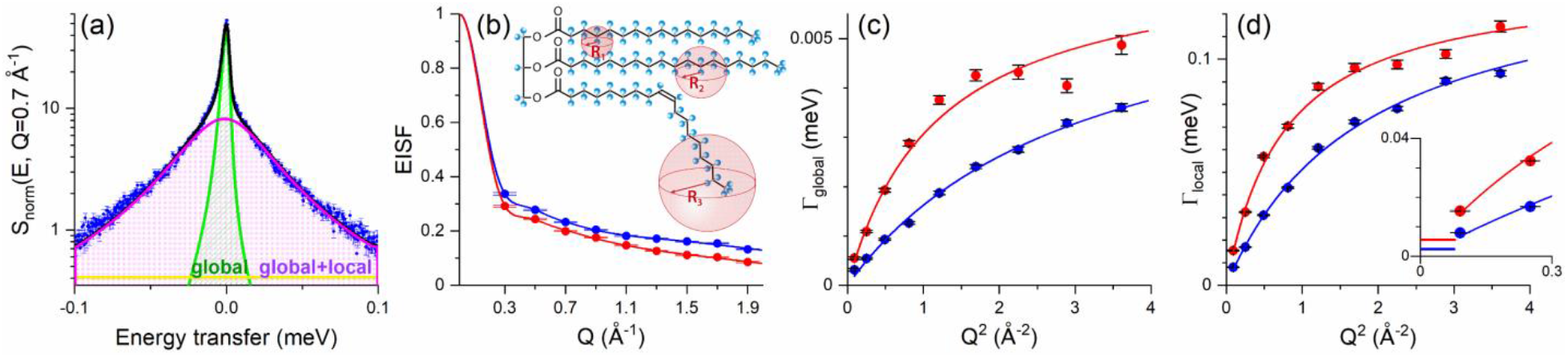
(a) Area-normalized QENS spectrum of the LDs at Q=0.7 Å^-1^ and T=280 K with the total fitting (black), convoluted Lorentzians for global (green) and total (magenta) dynamics, and the background (yellow). (b) The elastic incoherent structure factor (EISF) of the lipid chain diffusion in the yeast droplets at 280 K (blue) and 310 K (red). Solid lines represent fitting according to the diffusion law in three spheres; inset: a sketch of the TAG with assigned spheres to the hydrocarbon chains. Γ(Q) as a function of Q^2^ for global (c) and local (d) lipid dynamics in the droplet at 280 K (blue) and 310 K (red). The fitting follows the jump-diffusion law; inset: calculated constant Γ(Q) when boundary of the largest volume for local lipid dynamics is approached.

Fig. 3(b) shows the EISF of the lipid chain motion at the two extreme temperatures being best fitted (lowest chi-squared) by the function describing diffusion inside three spheres of different radii:

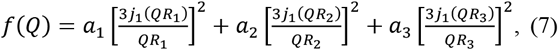

where *a*_3_ *=* 1 − *a*_1_ − *a*_2_, *a*_*i*_ is a spectral fraction of diffusion, *R*_*i*_ is a sphere radius, and *j*_1_ is the first-order spherical Bessel function. The obtained parameters are listed in Table I.

**TABLE I.**
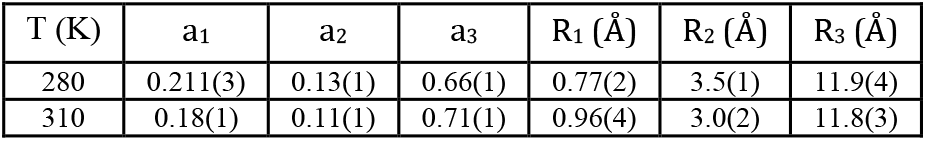
Relative contributions *a*_*i*_ and radii *R*_*i*_ derived from the fit to the EISF.

The resulting three radii are well distributed and are minimally affected by temperature. Thus, we define them as the small, medium, and large spheres for the local diffusion. The local motion of the fluid-phase DMPCs in the membrane was also described by the law for diffusion in a sphere. However, the number of spheres corresponded to the number of CH_2_ units in the lipid chain [22]. In that distribution the minimum and maximum radii, *R*_*min*_ and *R*_*max*_, were 0.04 and 5.15 Å, respectively. In the present case, the radii of small and medium spheres were comparable to those, except for the third sphere with a much larger radius. Having a much larger and temperature-independent volume for diffusion of the lipid chains, the yeast LDs were likely in a fluid phase at 280– 310 K. Moreover, structurally bent unsaturated lipids in the LDs are expected to cause less efficient packing of lipids that creates a space for their high diffusivity. A possible association of the spherical geometry with hydrocarbon chains of the TAG is depicted in the inset of Fig. 3(b).

How atoms diffuse, continuously or by discrete jumps, can be determined from the Q-dependence of the spectral width of the QENS signal. If the spectral width depends linearly on *Q*^2^, the atoms diffuse continuously. In the present case, the Lorentzian HWHM of both global and local lipid motions deviated from linearity and reached a plateau at higher *Q*^2^ (Fig. 3(c, d)). Such dependence is typical for a diffusion via some discrete translations and approximated by a jump-diffusion law. Here, we applied the function with an exponential distribution of jump lengths [31]:

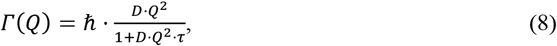

where *D* is the diffusion coefficient and *τ* is the residence time, i.e., the time when atoms oscillate between jumps. A jump-diffusion process is usually observed in the case of strongly interacting atoms. The fitting parameters for the lipid diffusion at the two extreme temperatures are listed in Table II. The diffusion coefficients of global and local motions differ significantly in accordance with the associated motion and are temperature dependent.

**TABLE II.**
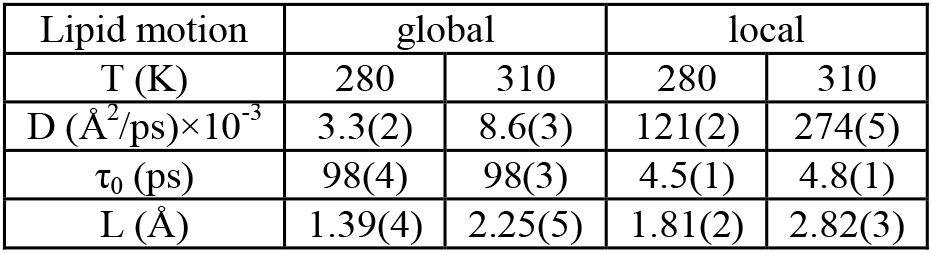
Fitting parameters of the jump-diffusion law for global and local motions of lipids in the droplet. *L* is the jump-length calculated as 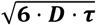.

In the case of spatially unrestricted diffusion, HWHM approaches zero at infinitely small Q (large length scale). In the case of diffusion in a limited space, HWHM approaches a plateau at Q→0 when boundary of the limited space is reached. According to the law of diffusion in a sphere [30], HWHM is expected to be constant and equal to 4.33ℏ*D* / *R*^2^ in the Q range from 0 up to 3.3 / *R*. With the largest sphere for the lipid chain motion, the biggest Q value is ∼0.3 Å^-1^ which is slightly smaller than the first Q value accessed in the experiment. Thus, the plateau in HWHM could not be identified experimentally. The expected constant values of HWHM for the lipid chain diffusion were calculated at both temperatures and marked in the inset of Fig. 3(d). The QENS describing local dynamics consists of three contributions with dominating one for the diffusion in the large sphere. The diffusion parameters for local dynamics are assumed the same in all spheres.

The diffusion coefficient of global lipid motion in the LDs at 310 K is 8.6(3)×10^−3^ Å^2^/ps, and is comparable to 7.7(3)×10^−3^ Å^2^/ps in the DMPC bilayers [22]. At 280 K the diffusion of whole lipid decreases to 3.3(2)×10^−3^ Å^2^/ps (∼2.5 times), which indicates still high lipid mobility in the droplet and an additional proof of their fluid phase. At this low temperature the DMPCs were in a gel phase and their diffusion reduced to 0.7(1)×10^−3^ Å^2^/ps (∼one order of magnitude less). From 280 to 310 K the diffusion coefficient of local lipid motion in the LDs increased from 121(2)×10^−3^ to 274(5)×10^−3^ Å^2^/ps. To compare, the highest coefficient of local diffusion in the fluid-phase DMPCs was 79(7)×10^−3^ Å^2^/ps [22]. The higher coefficients in the LDs are due to the larger sphere (R_3_=11.8 Å) for the local diffusion than in the bilayers (R_max_=5.15 Å).

The QENS study of human low-density lipoprotein [32], which has a similar structure with the LDs, reports the diffusion coefficients of translational and fast lipid motions being 11(2)×10^−3^ and 210(10)×10^−3^ Å^2^/ps, respectively, at 310 K that are in a good agreement with the LDs’ coefficients.

The diffusion parameters for lipids in the yeast droplet were reproducible as followed from the analysis of our second experiment with annular sample holder (the details on both measurements are reported in the SM [25]). Specifically, at 310 K it revealed the coefficients of 8.2(2)×10^−3^ Å^2^/ps for global and 237(5)×10^−3^ Å^2^/ps for local lipid diffusion, which reduced to 3.8(2)×10^−3^ Å^2^/ps and 114(2)×10^−3^ Å^2^/ps, respectively, at 280 K. The sphere radii for local lipid diffusion appeared the same at both temperatures as in the analysis of our first experiment.

In summary, the lack of a phase transition and the high diffusion coefficients of lipids in the droplets at 280–310 K provide insight into why a fluid phase of the LDs is maintained by the yeast in this temperature range. The physical data of the yeast LDs correlate with their chemical composition made of half of the unsaturated lipids contributing to a low packing ability of the lipids. It could be inferred that yeast *Cryptococcus curvatus* adopts the strategy of limiting the saturated-to-unsaturated lipid ratio in the LDs to ensure the fluid state of the droplets through the broad temperature range, which yeast may encounter in its natural environment, even at the expense of lowering lipid packing ability and energy storage efficiency. For future biofuel applications, where yeast is maintained at a stable temperature, such as in a biorefinery, and potential solidification of the LDs can be avoided, the goal should be to modify the microorganism to make it produce LDs with a higher saturated-to-unsaturated lipid ratio. Such advance would enable to obtain more energy from LDs for biofuels.

Our experiments revealed intrinsic lipid dynamics characterized by diffusion parameters in natural LDs probed in intact yeast cells. These findings represent an advance in lipid science, which heretofore has studied mostly synthetic lipid systems in aqueous solution. Moreover, by proving the reproducibility of dynamic data, we validated the suitability of the QENS technique for studies of biosystems *in vivo* to explore intrinsic microscopic dynamics and phase behavior.

## Acknowledgements

We thank Kevin Weiss and Frederick Heberle for the technical assistance with sample preparation and GC/MS measurements, James Shaw for the assistance with analysis of a 3D microscopic image of our samples using software IMARIS, and Deborah Counce for the valuable comments on text editing.

The neutron scattering experiments were sponsored by the Spallation Neutron Source (SNS), the U.S. Department of Energy (DOE) Office of Science User Facility operated by the Oak Ridge National Laboratory. The sample preparation was done at the SNS’s Biology Laboratory sponsored by the SNS and the U.S. DOE Office of Science Biological and Environmental Research.

