## Supplemental Material for "Dynamics of lipids in the yeast lipid droplets"

### Sample preparation

The appropriate growth medium for the yeast cells to accumulate LDs is a soluble starch 5% (SS5) having overabundant carbon content. Due to a deficiency of non-carbon nutrients in the medium, cell proliferation in the SS5 is slow (~120 h). The growth kinetics of the cells was evaluated by measuring the optical density (OD) of the cell culture at $\lambda$=600 nm using an Ultrospec 10 cell density meter from Amersham Biosciences. The accumulation of LDs in the cells was monitored periodically by fluorescence measurement at a wavelength of $\lambda$=660 nm using a Molecular Devices SpectraMax i3® (Fig. 1). The cell proliferation in a standard yeast malt extract (YME) medium having enough non-carbon nutrients, is fast. The maximum OD of the yeast culture in the YME is reached in ~18 h and is 10 times higher than the maximum OD reached in the SS5. To have a high cell density in a short time, the yeast *Cryptococcus curvatus* ATCC20509 was cultivated in 100 ml of $H_2O$-based YME in a 1-liter baffled flask shaken for air circulation at a preferred speed of 250 rpm and at T=303 K. Upon reaching the maximum OD in the YME, a 10$^{th}$ part of the cell pellet was transferred to the same amount of $H_2O$-based SS5 to trigger the accumulation of LDs. After 38 h, the yeast cells in the SS5 had formed LDs that were detectable under microscope (Fig. 2). At this stage, the cell culture was centrifuged at 4000×g for 10 min, the supernatant was removed, and the cell pellet was loaded into the sample holder for immediate QENS measurement. Keeping the cells in the SS5 for a longer time made the LDs bigger. However, large droplets overfilled the cells. Centrifugation of the yeast culture with the biggest LDs resulted in cell disruption and the release of the LDs. To obtain whole cells carrying LDs, the cells were harvested after 38 h. For a reference sample, i.e., cells without LDs (later, cells), the yeast culture cultivated in the YME was used. The mass of each sample, i.e., pellet of cells and cells with LDs, was ~0.5 g. The recipe for the YME and SS5 is reported in Table I. The sample preparation doesn't involve deuteration. Obtaining deuterated cells with hydrogenated LDs is uncertain approach because LD accumulation is not an isolated process and hydrogenated glucose will be used in both cell and LD growth.

| Ingredient | YME | SS5 |
|---|---|---|
| Peptone | 0.5% | |
| Malt extract | 0.3% | |
| Yeast extract | 0.3% | 0.01% |
| Glucose | 1% | 5.263% |
| $MgSO_4$ | | 0.5% |
| NaCl | | 0.01% |
| $CaCl_2$ | | 0.01% |

**TABLE I.** The recipe for the YME and SS5 growth media in 100 ml of $H_2O$.

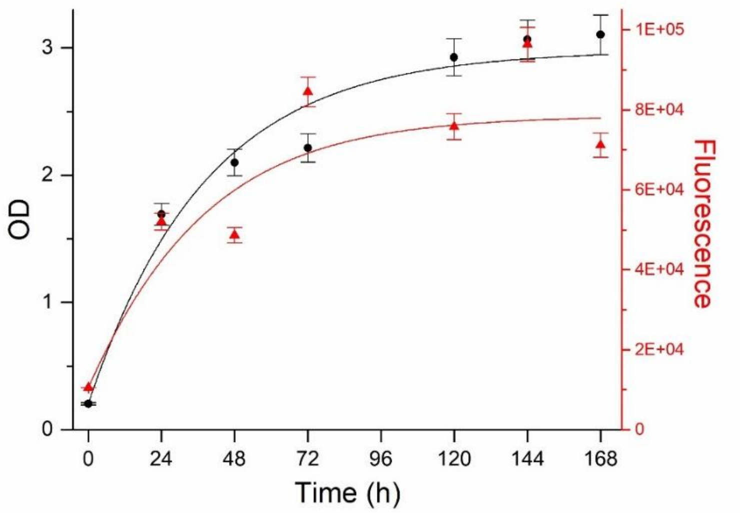

**FIG. 1.** The kinetics of the yeast cells and their LD formation in the SS5.

The lipid composition of the yeast droplets was determined using gas chromatography/mass spectrometry (GC/MS). The LDs were isolated from the cells following the protocol reported in [1]. The fatty acids of the yeast LDs were determined by fatty acid methyl ester analysis, as described in [2]. Analysis of the GC/MS spectrum of the isolated yeast droplets revealed the major part, 46%, consisted of oleic acid (C18:1 cis-9); 30 and 15% consisted of palmitic (C16:0) and stearic (C18:0) acids, respectively; and the remaining 9% was represented by other saturated and unsaturated acids (Fig. 2). Thus, almost equal proportions of monounsaturated (one double carbon bond; thus, structurally bent) and saturated (all single carbon bonds; thus, structurally straight) lipids made up the yeast droplet.

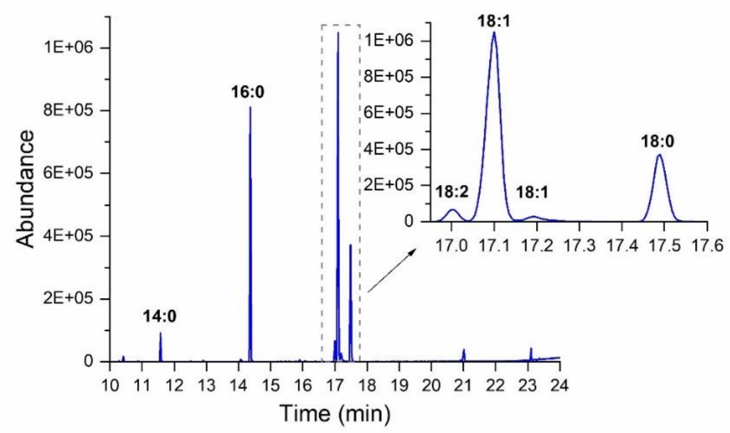

| C:D | % of total | Type |
|---|---|---|
| 14:0 | 3 | Sat. |
| 16:0 | 30 | Sat. |
| 18:2 | 3 | Unsat. |
| 18:1 | 46 | Unsat. |
| 18:0 | 15 | Sat. |
| other | 3 | Sat. |
| | 51 | Sat. |
| | 49 | Unsat. |

**FIG. 2.** The GC/MS spectrum of the isolated yeast LDs.

The volume ratio of the LDs to their cells was estimated using a 3D confocal microscope. For this purpose, the yeast culture was stained with the lipid-specific dye Nile Red (binds to lipid) [3] and the cell membrane–specific dye Calcofluor White (binds to chitin) [4]. The 3D images of the stained cells with LDs were obtained using a Zeiss confocal microscope with an oil-immersed 63X objective. One, two, or a few LDs were detected in the cells. Using IMARIS 3D image analysis software [5], the surfaces of the LDs and the cells were generated based on the intensity of the dyes (Fig. 3 (left)). The volumes of the surfaces were also quantified with IMARIS. The average LD-to-cell(s) volume ratio was ~43% (Fig. 3 (right)). Thus, the remaining 57% of the cellular volume was ascribed to the cytoplasm and other cellular organelles.

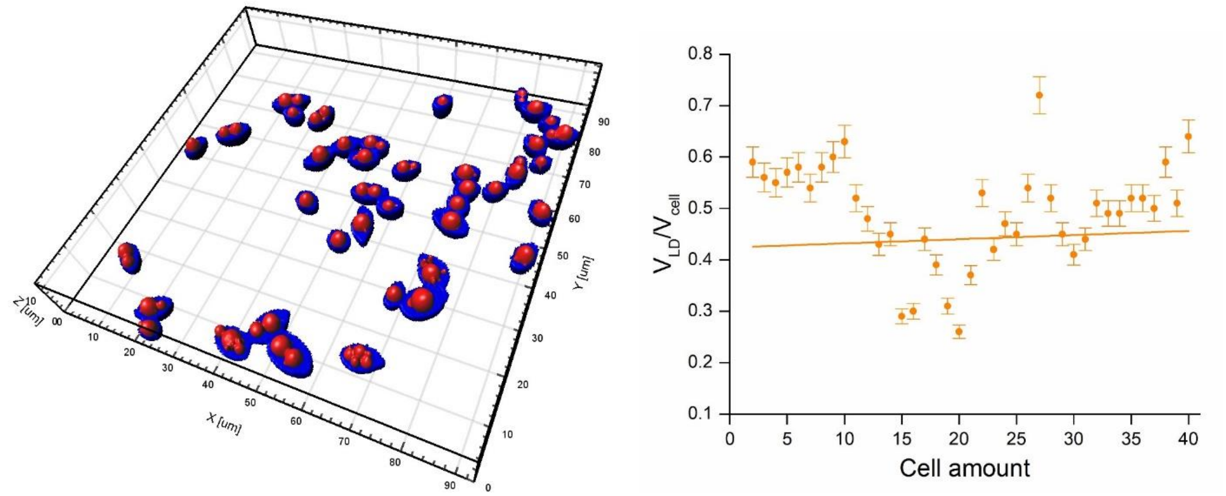

**FIG. 3.** (left) A 3D microscopic image of yeast cells. The cell surfaces are shown in blue color, the LDs are highlighted in red. (right) The LD(s)-cell volume ratio estimated in IMARIS.

### QENS

The experiment was performed twice in separate events to check the reproducibility of the data obtained for the yeast LDs. Aluminum sample holders of different geometries were used in the experiments. For the first experiment, a rectangular holder of 50×30 $mm^2$ (h×w) was used; and for the second, an annular holder of 54×29 $mm^2$ (h×diameter) was used, both having a 0.25 mm gap for the sample. The neutron beam was limited to 30×30 $mm^2$ (h×w) and directed perpendicular to the sample. The incident bandwidth choppers were operated at 60 Hz, and the wavelength of the neutrons reflected from the Si(111) analyzers at an 88° Bragg angle was 6.267 Å. The instrument energy resolution averaged over all scattering angles is typically 0.0034 meV, which is the full width at half maximum (FWHM). The energy transfer bandwidth was ±0.1 meV, which, together with the spectrometer energy resolution, defined the accessible time scale of 6–400 picoseconds. The data obtained at the scattering angles corresponding to the wavevector transfer Q of 0.3–1.9 $Å^{-1}$ defined the accessible length scale of 3–20 Å.

The QENS raw data, initially in neutron time-of-flight format, were converted to the dynamic structure factor $S(E,Q)$ and corrected for the detector efficiency by normalization to a vanadium standard. Then $S(E,Q)$ spectra were grouped at every 0.0004 meV in the $E$ and at 0.2 $Å^{-1}$ in the $Q$ to improve the statistics. The QENS data on the microscopic dynamics were obtained at 280 and 310 K for a few hours. The data on the phase behavior were obtained from 280 to 310 K every 0.5 K with a scan rate of 0.4 K/min, and then integrated over the instrument energy resolution and summed over the total Q range to derive the total incoherent elastic intensity at each temperature. The instrument energy resolution was measured on the cells having LDs at 10 K, where all measurable dynamics was frozen out. The instrument energy resolution was used in numerical convolution with the model function (equation 6 in the main text) for the spectra fitting. Because of the convolution, the sample dynamics (HWHM) is resolvable at the energies below the instrument resolution. Typically, accumulating scattering signal from the sample for a few hours allows to resolve dynamics of the sample down to 10-20% of the instrument resolution's HWHM. The empty holder was measured to subtract its contribution from the sample spectra.

To derive the QENS signal of the LDs, the cell contribution had to be subtracted from the spectra of cells with LDs. To estimate the cell contribution, the spectra of cells were divided by 2.7, which was obtained by the following procedure. A 3D surface modeling in the IMARIS software obtained the volume fractions of 43% for the LD and 57% for its cell (Fig. 3). Measuring samples of equal mass meant having more cells than cells with LD. Based on lyophilization of both samples, the mass fractions of 63% for the LD and 37% for its cell were determined. Through the density equation, the volume fraction of the cells to that of the cells with LDs produced the factor of 2.7. The data reduction and analysis were conducted in the MANTID software [6].

The neutron scattering experiment was carried out at the Spallation Neutron Source (SNS) located at Oak Ridge National Laboratory (ORNL). The QENS measurements of the yeast pellets were performed at the backscattering time-of-flight spectrometer BASIS [7]. This instrument resolves microscopic dynamics in the sample in time and space.

[1] Y. Ding, S. Zhang, L. Yang, H. Na, P. Zhang, H. Zhang, Y. Wang, Y. Chen, J. Yu, C. Huo, S. Xu, M. Garaiova, Y. Cong, and P. Liu, Nat. Protoc. **8**, 43 (2013).

[2] F. A. Heberle, D. Marquardt, M. Doktorova, B. Geier, R. F. Standaert, P. Heftberger, B. Kollmitzer, J. D. Nickels, R. A. Dick, G. W. Feigenson, J. Katsaras, E. London, and G. Pabst, Langmuir **32**, 5195 (2016).

[3] S. D. Greenspan, P.; Mayer, E. P.; Fowler, J. Cell Biol. **100**, 965 (1985).

[4] G. J. Hageage and B. J. Harrington, Lab. Med. **15**, 109 (1984).

[5] http://www.bitplane.com/

[6] http://dx.doi.org/10.5286/SOFTWARE/MANTID.

[7] E. Mamontov and K. W. Herwig, Rev. Sci. Instrum. **82**, (2011).

### Additional equations on normalization of $S(E,Q)$

The total dynamic structure factor including definition of global and local structure factors:

$$
\begin{aligned}
S(E,Q) &= A \cdot L_{global}(E) \otimes B \cdot \left[f(Q) \cdot \delta(E) + \left(1 - f(Q)\right) \cdot L_{local}(E)\right] \cdot e^{-\langle u^2(T)\rangle \cdot Q^2} \\
&= A \cdot B \cdot e^{-\langle u^2(T)\rangle \cdot Q^2} \cdot \left[f(Q) \cdot L_{global}(E) + (1 - f(Q)) \cdot L_{total}(E)\right]
\end{aligned}
$$

As the Lorentzian functions are area-normalized, the total dynamic structure factor should be normalized as well:

$$
\begin{aligned}
\frac{S(E,Q)}{\int_{-\infty}^{+\infty} S(E,Q)dE} &= \frac{A \cdot B \cdot e^{-\langle u^2(T)\rangle \cdot Q^2} \cdot \left[f(Q) \cdot L_{global}(E) + (1 - f(Q)) \cdot L_{total}(E)\right]}{A \cdot B \cdot e^{-\langle u^2(T)\rangle \cdot Q^2} \cdot \int_{-\infty}^{+\infty}\left[f(Q) \cdot L_{global}(E) + (1 - f(Q)) \cdot L_{total}(E)\right] dE} \\
&= f(Q) \cdot L_{global}(E) + (1 - f(Q)) \cdot L_{total}(E)
\end{aligned}
$$

The normalized total dynamic structure factor including definition of the Lorentzian function:

$$
S_{norm}(E,Q) = f(Q) \cdot \frac{\Gamma_{global}(Q)}{\pi\left({\Gamma_{global}}^2(Q) + E^2\right)} + (1 - f(Q)) \cdot \frac{\Gamma_{global}(Q) + \Gamma_{local}(Q)}{\pi\left[(\Gamma_{global}(Q) + \Gamma_{local}(Q))^2 + E^2\right]}
$$
